# MAP9/MAPH-9 supports axonemal microtubule doublets and modulates motor movement

**DOI:** 10.1101/2023.02.23.529616

**Authors:** Michael V. Tran, James W. Ferguson, Lauren E. Cote, Daria Khuntsariya, Richard D. Fetter, Jennifer T. Wang, Stephen R. Wellard, Maria D. Sallee, Mariya Genova, Sani Eskinazi, Maria M. Magiera, Carsten Janke, Tim Stearns, Zdenek Lansky, Kang Shen, Jérémy Magescas, Jessica L. Feldman

**Affiliations:** Department of Biology, Stanford University, Stanford, CA 94305, USA; Institute of Biotechnology, Czech Academy of Sciences, BIOCEV, Prague West, Czech Republic; Howard Hughes Medical Institute, Department of Biology, Stanford University, Stanford, CA, USA; Department of Biology, Washington University in St. Louis, St. Louis, Missouri, USA; Institut Curie, Université PSL, CNRS UMR3348, Orsay, France; Université Paris-Saclay, CNRS UMR3348, Orsay, France; Department of Genetics, Stanford University School of Medicine, Stanford, CA 94305, USA

**Keywords:** Microtubule, Microtubule doublet, microtubule-associated protein, MAP9, polyglutamylation, dynein, kinesin, microtubule, cilia, axoneme, *C. elegans*

## Abstract

Microtubule doublets (MTDs) are a well conserved compound microtubule structure found primarily in cilia. However, the mechanisms by which MTDs form and are maintained *in vivo* remain poorly understood. Here, we characterize microtubule-associated protein 9 (MAP9) as a novel MTD-associated protein. We demonstrate that *C. elegans* MAPH-9, a MAP9 homolog, is present during MTD assembly and localizes exclusively to MTDs, a preference that is in part mediated by tubulin polyglutamylation. Loss of MAPH-9 caused ultrastructural MTD defects, dysregulated axonemal motor velocity, and perturbed cilia function. As we found that the mammalian ortholog MAP9 localized to axonemes in cultured mammalian cells and mouse tissues, we propose that MAP9/MAPH-9 plays a conserved role in supporting the structure of axonemal MTDs and regulating ciliary motors.

## Introduction

Microtubules are intracellular polymers involved in essential cell processes ranging from cell division to maintaining cell structure. Microtubules are made up of α- and β-tubulin heterodimers that associate end-to-end to form protofilaments that associate laterally to form a single hollow cylinder. However, in specialized cellular appendages called cilia, microtubules are instead found in a figure-8 like doublet structure composed of a single microtubule (the A-tubule) attached to an incomplete microtubule (the B-tubule) (Figure 1B). Most cilia have nine microtubule doublets (MTDs) arranged in a radially symmetric array to form the axoneme. The axoneme provides the tracks for intraflagellar transport (IFT), the process by which molecular motors (anterograde kinesin and retrograde dynein) transport cargo to build and maintain the cilium and support cilia function.

**Figure 1.**
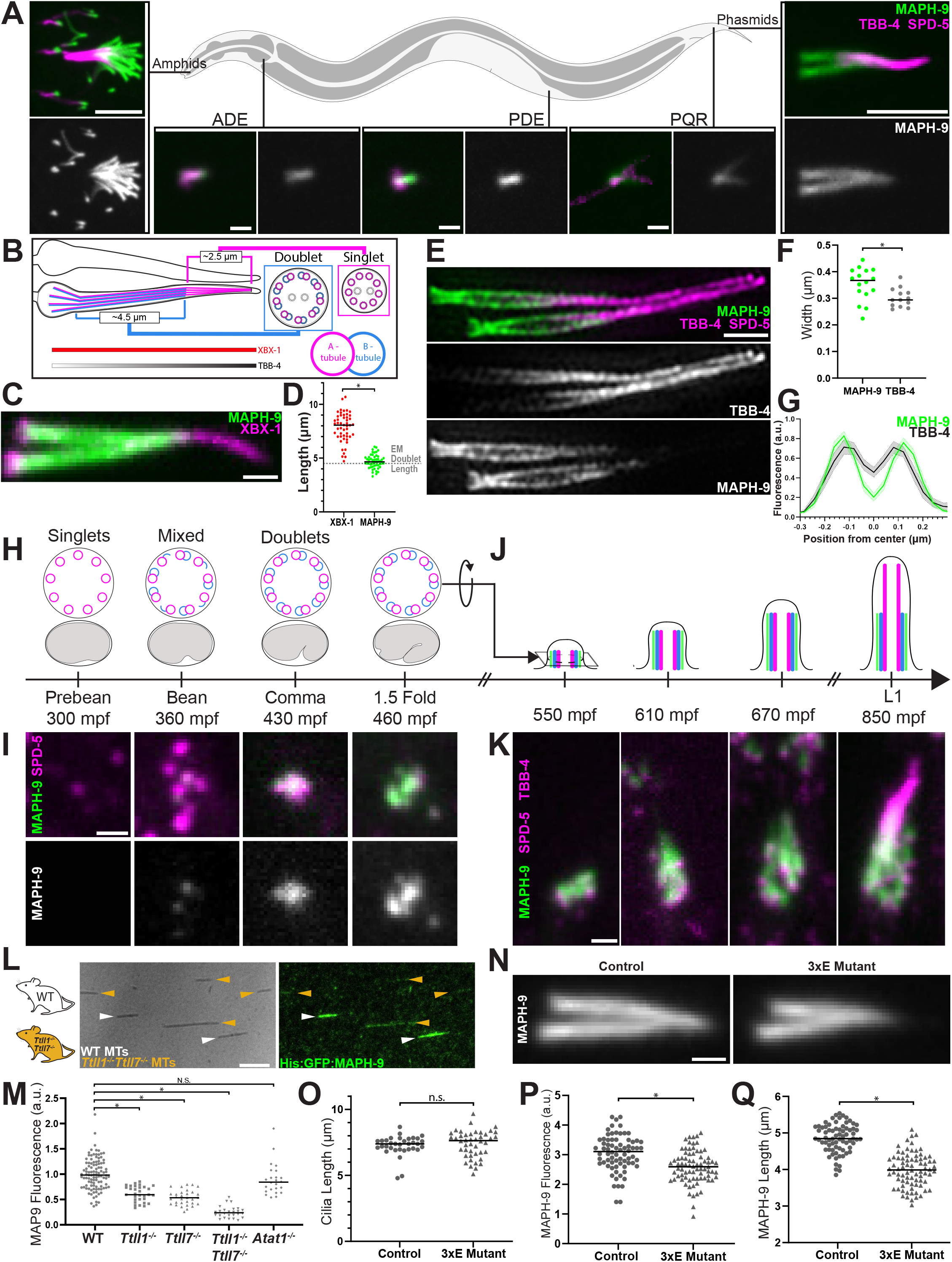
MAPH-9 localizes to microtubule doublets and preferentially binds polyglutamylated microtubules. (A) Spinning disk confocal imaging of endogenously tagged GFP::MAPH-9 (green), TBB-4::RFP, and RFP::SPD-5 (magenta) in indicated ciliated sensory neurons in adult worms. TBB-4/SPD-5 brightness scaled differently in PQR. Scale bars: Amphids and Phasmids, 5 μm; ADE, PDE, PQR, 1 μm. (B) Schematic depicting the cilia of the phasmid neurons within one axoneme highlighted and its cross section in the doublet or singlet region with the A-tubule (magenta) and B-tubule (blue) indicated. XBX-1 and TBB-4 localize to the length of the axoneme. (C) Localization of endogenous MAPH-9 (green) and XBX-1 (magenta) in adult phasmid cilia. Scale bar, 1 μm. (D) Quantification of protein localization length (μm) from ciliary base marked by XBX-1. Dashed horizontal line is length of doublet region in phasmid cilia (∼4.5 μm). XBX-1: 7.86±1.33 n=50; MAPH-9: 4.69±0.63 n=50; P<0.0001. (E) 3D structured illumination (3D SIM) imaging of endogenously tagged MAPH-9 (green) and TBB-4/SPD-5 (magenta) in adult phasmid cilia. Scale bar, 1 μm. (F) Quantification of width (μm) of MAPH-9 and TBB-4 fluorescence from 3D SIM imaging measured perpendicular to length of axoneme in the middle segment, i.e. doublet region. TBB-4: 0.30±0.06 n=13; MAPH-9: 0.35±0.03 n=16; P<0.05. (G) Averaged 3D SIM line profiles of MAPH-9 (green) and TBB-4 (black) fluorescence (a.u.) perpendicular to length of axoneme in the middle segment. (H) Schematic depicting the timeline of development of microtubule doublets at centrioles (top, cross-sectional view) in amphid neurons during *C. elegans* embryonic development (bottom, side view) at indicated minutes post-fertilization (m.p.f.) (Nechipurenko et al., 2017). (I) Localization of endogenously tagged MAPH-9 (green) and SPD-5 (magenta) in amphid neurons through embryonic development. Scale bar, 1 μm. (J) Longitudinal view schematic depicting A-tubule (magenta), B-tubule (blue), and MAPH-9 (green) localization during ciliogenesis in the embryo at indicated m.p.f. (K) Localization of endogenously tagged MAPH-9 (green) and SPD-5/TBB-4 (magenta) during ciliogenesis. Scale bar, 1 μm. (L) Total Internal Reflection Fluorescence microscopy images of: Left: microtubules assembled from tubulin from wildtype (WT) mice (white arrowheads) or *Ttll1*^−/−^ and *Ttll7*^−/−^ mice (orange arrowheads); Right: MAPH-9 localization. Scale bar, 5 μm. (M) Quantification of normalized fluorescence intensity (a.u.) of MAPH-9 localized to microtubules assembled from tubulin extracted from WT, *Ttll1*^−/−^, *Ttll7*^−/−^, *Ttll1*^−/−^ and *Ttll7*^−/−^, or *Atat1*^−/−^ mice. Control: 1.0±0.31 n=105; *Ttll1*^−/−^: 0.61±0.15 n=31 P<0.0001; *Ttll7*^−/−^: 0.54±0.15 n=31 P<0.0001; *Ttll1*^−/−^ *Ttll7*^−/−^: 0.25±0.11 n=26 P<0.0001; *Atat1*^*−/−*^: 0.89±.31 n=24 P=0.15. P-values compared to control. (N) Localization of MAPH-9 in phasmid cilia in control and *ttll-4(tm3310), ttll-5(tm4059), ttll-11(tm3360)* polyglutamylation mutant (3xE mutant). Scale bar, 1 μm. (O) Length (μm) of cilia measured from the base of MAPH-9 (anterior end) to the posterior end of DiI dye filling. Control: 7.25±0.75 n=34; 3xE mutant: 7.39±1.01 n=44; P=0.522. (P) Normalized mean fluorescence of MAPH-9 per unit length of cilia (a.u.). Control: 3.05±0.61 n=72; 3xE mutant: 2.59±0.55 n=77; P<0.0001. (Q) Length (μm) of MAPH-9 localization. Control: 4.9±0.38 n=72; 3xE mutant: 4.0±0.45 n=78; P<0.0001. Values presented are mean±SD. P-values are calculated by Welch’s t-test. Graphs present individual data points with horizontal bar representing the median.

MTD formation *in vitro* can be driven by subtilisin treatment which cleaves the C-terminal tails from α- and β-tubulin (Schmidt-Cernohorska et al., 2019), however, how MTD formation is stimulated *in vivo* is poorly understood. In the axoneme, a collection of microtubule inner proteins (MIPs) that bind to the inside of the microtubule protofilament like FAP52 and FAP20 in *Chlamydomonas* are thought to contribute to the MTD (Nicastro et al., 2006; Owa et al., 2019; Pigino et al., 2012; Sui & Downing, 2006; Yanagisawa et al., 2014). Depletion of ε-tubulin, δ- tubulin, centriolar protein HYLS-1, or mutation in *Arl13b* disrupts the formation of compound microtubules; however, ε-tubulin and δ-tubulin are not conserved in all species with cilia, HYLS-1 does not localize to axonemal MTDs, and Arl13b is a ciliary membrane-associated G-protein where it likely interacts indirectly with the axoneme, suggesting other factors are involved in MTD formation and maintenance (Caspary et al., 2007; Cevik et al., 2010; Dutcher et al., 2002; Mottier-Pavie & Megraw, 2009; Wang et al., 2017; Wei et al., 2016).

While MIPs and other factors contribute to MTDs, the identity and role of classic lattice binding microtubule-associated proteins (MAPs) at MTDs remain largely unknown (Bodakuntla et al., 2019; Conkar & Firat-Karalar, 2021). Although originally shown to localize to cytoplasmic microtubules and to play roles in the mitotic spindle (Saffin et al., 2005), the understudied MAP9/ASAP has been implicated in cilia-related processes: a deletion in *MAP9* enhances retinal degeneration in Miniature Long-Haired Dachshunds caused by an insertion in the gene for ciliary protein RPGRIP1 (Forman et al., 2016); *map9* depletion in zebrafish causes developmental defects associated with dysregulation of hedgehog signaling which takes place in primary cilia (Fontenille et al., 2014); and the *C. elegans* homolog MAPH-9 is specifically expressed in ciliated sensory neurons and localizes to cilia when overexpressed (Jensen et al., 2016; Venoux et al., 2008). Despite evidence for a ciliary role for MAP9, its precise function within this organelle has been largely unexplored.

We used *C. elegans* to explore the *in vivo* function of MAP9, finding a role for MAPH-9 specifically in cilia. Endogenous MAPH-9 localized exclusively to axonemal MTDs, distinguishing these unique structures from both cytoplasmic microtubules and microtubule singlets within the axoneme in part through a preference for polyglutamylated tubulin.

Furthermore, in the absence of MAPH-9 MTDs were either lost or perturbed, indicating a role for MAPH-9 in building and/or maintaining the MTD structure. Loss of MAPH-9 also modulated kinesin frequency and velocity and accelerated dynein velocity, indicating a specific role of MAPH-9 in modulating IFT motor movement. These cilia defects also had physiological consequences as *maph-9* null mutants had defects in mating, a process known to rely on cilia. As we found that MAP9 localization is conserved in mammalian motile and primary cilia, our work provides novel insight into the role of an understudied MAP in shaping the highly conserved MTDs of cilia.

## Results

### MAPH-9 is expressed in ciliated sensory neurons and specifically localizes to the doublet region of the axoneme

We explored the endogenous localization pattern of MAPH-9 in *C. elegans* by inserting GFP into the endogenous *maph-9* locus using CRISPR/Cas9 genome editing. Live imaging in young adult worms showed that GFP::MAPH-9 is exclusively expressed in all ciliated cells, which in *C. elegans* is a subset of sensory neurons that include groups in the head (amphids) and the tail (phasmids) (Figure 1A and Figure S1A). MAPH-9 colocalized with endogenously tagged β-tubulin/TBB-4::RFP, a tubulin isotype that specifically localizes to the ciliary axoneme (Hao et al., 2011; Nishida et al., 2021), and ∼CDK5RAP2/RFP::SPD-5, which marks the base of cilia (Figure 1A) (Garbrecht et al., 2021; Magescas et al., 2021). Intriguingly, although MAPH-9 localized to all cilia, its localization was restricted to the more proximal region of the axoneme.

To better understand this localization, we compared the length of MAPH-9 localization to the endogenously tagged dynein component D2LIC/XBX-1::RFP which localizes along the length of each cilium (Figure 1C). The length of MAPH-9 localization was significantly shorter than that of XBX-1 (XBX-1: 7.86±1.33 μm n=50; MAPH-9: 4.69±0.63 n=50; P<0.0001; Figure 1D) and notably the same as the MTD region of the axoneme (so-called middle segment in *C. elegans* cilia) as measured from serial electron microscopy reconstruction (EM, ∼4.5 μm; Figure 1B,1D) (Perkins et al., 1986; Ward et al., 1975), suggesting that MAPH-9 colocalizes with MTDs.

As MAPH-9 is a MAP and therefore predicted to bind directly to microtubules, we verified that it localizes to the axoneme using super-resolution microscopy. In longitudinal optical sections of the axoneme, GFP::MAPH-9 localized in two discrete bands (Figure 1E) with MAPH-9 localizing to a region that was slightly wider than TBB-4 (MAPH-9: 0.35±0.062 μm n=16; TBB-4: 0.30±0.036 μm n=13; p<0.05; Figure 1F). The width of GFP::MAPH-9 localization was significantly larger than the distance between axonemal microtubules measured from electron micrographs (Figure 1E-1G)(Doroquez et al., 2014; Perkins et al., 1986). These data suggest that MAPH-9 localizes around the membrane-facing side of the MTD region of the axoneme.

### MAPH-9 localizes to the centrioles concomitant with the appearance of microtubule doublets

To further test if MAPH-9 specifically recognizes MTDs, we took advantage of a unique aspect of ciliogenesis in *C. elegans*. Cilia are built from microtubule-based structures called centrioles that serve as a template to build the nine-fold arrangement of MTDs that constitute the axoneme. While centrioles in *C. elegans* are normally comprised of nine microtubule singlets, centrioles in ciliated cells build doublets *de novo* just prior to ciliogenesis (Figure 1H)(Nechipurenko et al., 2017). We therefore determined if MAPH-9 localizes to the centriole concomitant with the appearance of doublets, which occurs in the amphid neurons at “bean” stage of embryogenesis (∼360mpf, Figure 1H-1I). Each ciliated neuron has one centriole that stays associated with the nucleus and another that migrates out to the tip of the dendrite to build the axoneme (Li et al., 2017). GFP::MAPH-9 localized to both centrioles as indicated by its colocalization with the centrosome component SPD-5 beginning at bean stage (Figure 1H-1I). While the MAPH-9 signal that colocalized with the nuclear-associated SPD-5 punctum stayed consistently low, the other dendrite associated MAPH-9 punctum intensified steadily throughout embryogenesis (Figure S1B-1C), consistent with MAPH-9 recognizing MTDs.

To determine if MAPH-9 is loaded into preassembled axonemes or associates with microtubules as the axoneme is built, we tracked MAPH-9 localization throughout ciliogenesis in live embryos (Figure 1J-1K). As ciliogenesis occurs at the stage of development when embryos move rapidly, we imaged MAPH-9 along with TBB-4 and SPD-5 in embryos anesthetized with carbon dioxide gas. MAPH-9 co-localized with TBB-4 throughout ciliogenesis (Figure 1K and Figure S1D). Together these data indicate that MAPH-9 localizes to structures that contain MTDs as they are being built and suggests a role for MAPH-9 in building or maintaining this unique microtubule-based structure.

### MAPH-9 preferentially binds polyglutamylated microtubules

Given the ability of MAPH-9 to distinguish between doublet and singlet microtubules, we wanted to identify features of the MTD that could impart this specificity. MAP binding to microtubules is influenced by posttranslational modifications (PTMs) of tubulins (Gadadhar et al., 2017; Wloga et al., 2017), many of which occur in the axoneme (Mirvis et al., 2018). In particular, *C. elegans* cilia are highly polyglutamylated (Kimura et al., 2010), making it a particularly attractive PTM to investigate. To determine if the polyglutamylation state of microtubules impacts MAPH-9 binding, we tested the ability of purified MAPH-9 protein to bind *in vitro* to microtubules with and without polyglutamylation.

Tubulin with different levels and patterns of PTMs was purified from the brains of mice mutant for different tubulin-modifying enzymes: mutants for the polyglutamylases TTLL1 and/or TTLL7, which lack polyglutamylation on α- and/or β-tubulin, and for the acetyltransferase αTAT1, which lacks tubulin acetylation. Compared to fully posttranslationally modified microtubules from control mice, MAPH-9 showed a significantly reduced binding to microtubules lacking α- and/or β-tubulin polyglutamylation (Figure 1L-1M and Figure S1E). This binding preference was specific to polyglutamylation as modulation of microtubule acetylation had no impact on MAPH-9 binding (Figure 1M and Figure S1E). The observation that the lack of both TTLL1-catalyzed α-tubulin polyglutamylation and TTLL7-catalyzed β-tubulin polyglutamylation affected MAPH-9 binding, and that loss of both had an additive effect, suggested that polyglutamylation has a general, and not an enzyme-specific effect on MAPH-9-microtubule interactions. We thus assumed that polyglutamylation catalyzed by other enzymes, i.e. the enzymes expressed in *C. elegans*, could also impact MAPH-9 interactions with microtubules. To assess the role of polyglutamylation on MAPH-9 localization to MTDs *in vivo*, we compared the localization of MAPH-9 in worms lacking the glutamylases TTLL-4, TTLL-5, and TTLL-11, a condition shown to eliminate axonemal polyglutamylation in *C. elegans* (Chawla et al., 2016). In *ttll-4, ttll-5, ttll-11* triple mutants (3xE mutant), phasmid cilia length was unaffected (Figure 1O), however the region of MAPH-9 localization was significantly shorter than in control worms and there was less MAPH-9 per unit length (Figure 1N-1Q). Together, these data indicate that polyglutamylation attracts MAPH-9 localization to microtubules *in vivo* and *in vitro*.

### Loss of MAPH-9 causes ultrastructural microtubule doublet defects in the axoneme

To determine whether MAPH-9 plays a structural role at MTDs, we generated a *maph-9* deletion mutant (*maph-9(0)*, Figure S1F). EM analyses of the amphid and phasmid neurons in *maph-9(0)* mutants showed that many of the B-tubules of the MTDs lacked the stereotypical roundness found in control axonemes (Figure 2A-2C, Figure S2A; WT:0.85±0.038 n=132; *maph-9(0)*:0.81±0.69 n=210; p<0.0001). Although this squashed B-tubule phenotype appeared to be present variably across amphid and phasmid cilia, we also observed additional neuron specific phenotypes. In the ADL neuron, *maph-9* deletion led to an absence of MTDs in cilia even though both control and *maph-9(0)* mutant worms had 9 MTDs at the base of the axoneme (Figure 2D-2E). In control ADL axonemes, these 9 MTDs extended into the middle segment, progressed to incomplete doublets, and eventually tapered to singlet microtubules before terminating at variable lengths near the distal portion of the cilium (Figure 2D-2E). In contrast, *maph-9(0)* mutant ADL axonemes had a significantly shorter region of MTDs and incomplete doublets, with the middle segment of the axoneme being comprised mainly of singlet microtubules (Figure 2D-2E). We observed a more extreme loss of microtubules in the axoneme of ASI neurons in *maph-9(0)* mutants, with axonemes completely lacking MTDs despite having 9 MTDs at their base (Figure S2B-S2C). These ultrastructural defects in *maph-9(0)* mutants suggest that MAPH-9 promotes the assembly and/or stability of MTDs within the axoneme.

**Figure 2.**
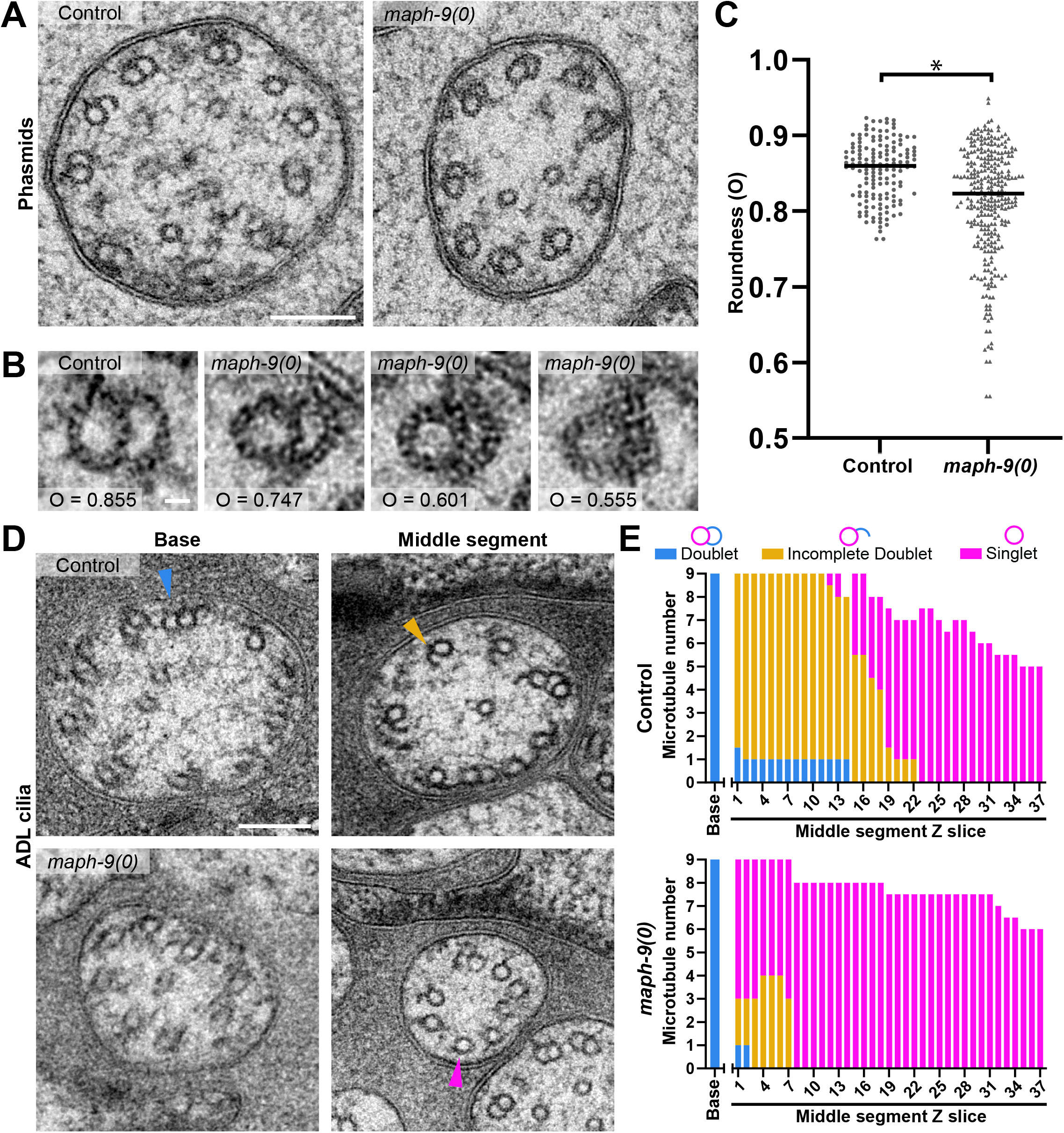
Loss of *MAPH-9 causes ultrastructural microtubule doublet defects in the axoneme*. (A) Representative EM images of cross sections through adult phasmid neurons of indicated genotypes. Scale bar, 100 nm. (B) Examples of individual doublets with genotype and roundness measurement (O) indicated. Scale bar, 10 nm. (C) Roundness measurement (O) of B-tubule in control and *maph-9(0)* mutants in phasmid neurons. Control: 0.85±0.04, n=3 worms, 148 sections; *maph-9(0)*: 0.81±0.07, n=4 worms, 305 sections; P<0.001. Mean±SD. P values are calculated by Welch’s t-test. (D) Representative EM images of amphid neuron ADL axoneme in control and *maph-9(0)* mutant worms. Arrowheads indicate representative doublet (blue), incomplete doublet (orange), and singlet (magenta) axonemal microtubules. Scale bar, 100 nm. (E) Quantification of doublet, incomplete doublets, and singlet microtubules in ADL axonemes. Measurement at base (left) and then in sections in the middle segment going toward the distal segment (right).

### MAPH-9 modulates motor activity and controls cilia function

MAPs are known to modulate the ability of molecular motors to bind to and walk on microtubules, potentially generating a code that directs traffic on the lattice (Bodakuntla et al., 2019; Gadadhar et al., 2017; Yu et al., 2015). *in vitro*, MAP9 inhibits microtubule binding of kinesin-1 and of dynein via dynactin while also promoting kinesin-3 activity (Monroy et al., 2020). However, the assembly and maintenance of cilia relies on IFT controlled by anterograde kinesin-2 motors and retrograde dynein motors that lack dynactin. *C. elegans* cilia use two semi-redundant kinesin-2 anterograde motors; the heterotrimeric kinesin-II, composed of KLP-11, KLP-20 and KAP-1, and the homodimeric OSM-3; with both motors walking on the doublet microtubules and only OSM-3 walking on the singlet microtubules at the distal portion of the axoneme (Figure 3A) (Prevo et al., 2015). OSM-3 is sufficient to build a full length axoneme in the absence of kinesin-II, but only loss of both kinesin-II and OSM-3 completely abrogates ciliogenesis (Evans et al., 2006). To understand if MAPH-9 works in concert with these motors to build or maintain cilia, we measured axoneme length in mutants lacking MAPH-9 and/or anterograde motor activity. *maph-9(0)* mutants had full length cilia (Figure 3B, Figure S1G-S1I). Similarly, loss of *maph-9* in an *osm-3* null mutant background (*osm-3(0))* did not further impact cilia length compared to an *osm-3(0)* mutant alone, indicating that kinesin-II still functions in the absence of MAPH-9 (Figure 3B, Figure S1I). In contrast, double mutants with both *maph-9(0)* and the essential kinesin-II component *kap-1* null mutant (*kap-1(0)*) showed a significant decrease in axoneme length compared to *kap-1(0)* mutants alone (Figure 3B and Figure S1I). These results suggest that MAPH-9 affects the function of OSM-3 more than that of kinesin-II or that OSM-3 is more sensitive to the loss of MAPH-9 when it is the only anterograde motor.

**Figure 3.**
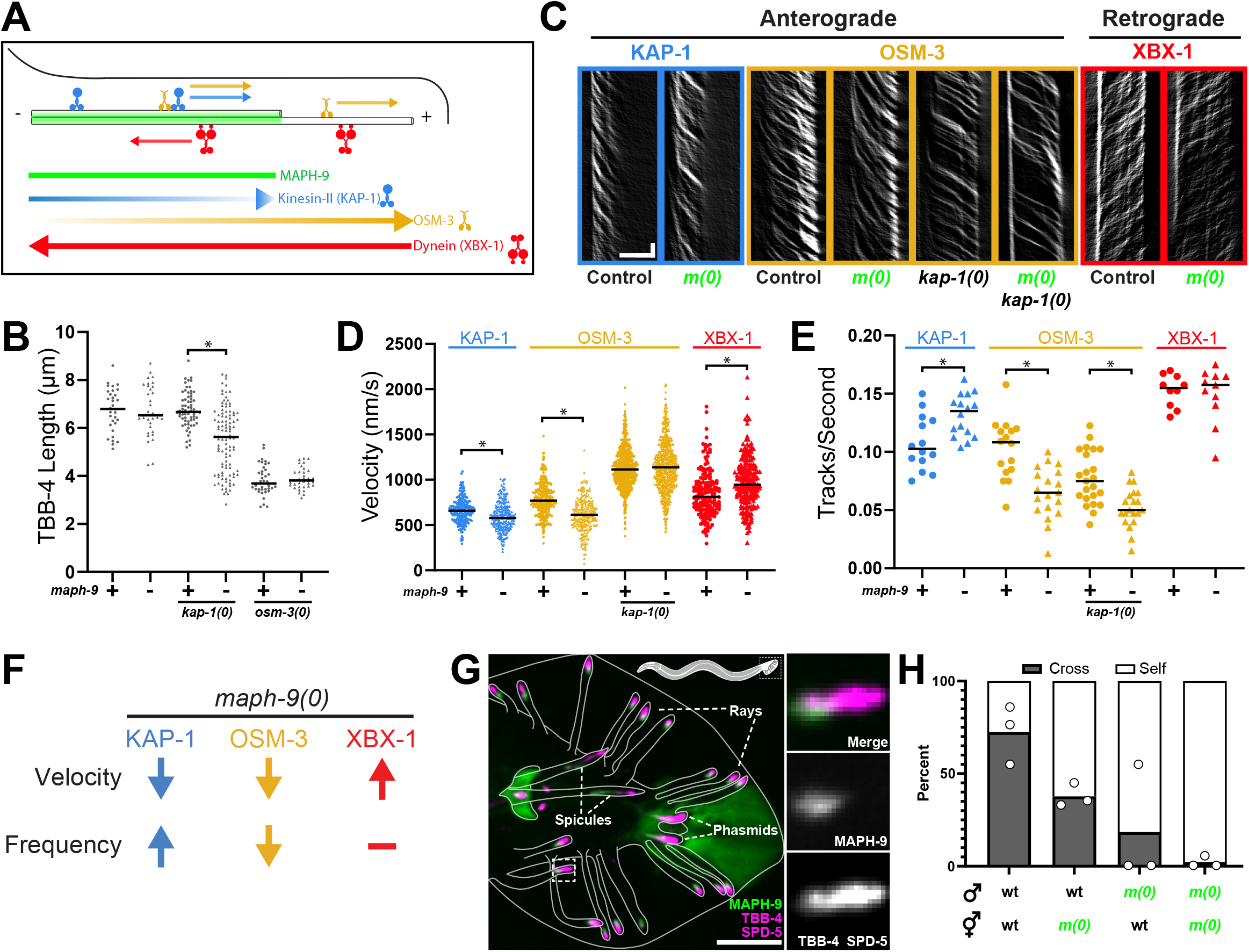
MAPH-9 modulates motor speed and controls cilia function. (A) Schematic depicting ciliary motors on the axoneme. Colored bars and arrows depict localization and direction of movement of proteins (anterograde to the right, retrograde to the left) in *C. elegans* cilia. (B) Quantification of length (μm) of TBB-4 fluorescence. Control: 6.75±0.87 n=32. *maph-9(0)*:6.74±1.06 n=32. *osm-3(0)*: 3.82±0.61 n=38. *osm-3(0)*; *maph-9(0)*:3.88±0.42 n=36. *kap-1(0)*:6.73±0.78 n=66. *kap-1(0)*; *maph-9(0)*:5.51±1.34 n=106. (C) Representative kymographs of KAP-1, OSM-3, and XBX-1 in control or mutant backgrounds. Scale bar, 5 μm (x); 1s (y). (D) Quantification of motor velocity (nm/s). KAP-1::GFP [Control: 660.7±134.8 n=240. *maph-9(0)*: 592.5±157.1 n=225]; OSM-3::GFP [Control: 781.9±181.7 n=319; *maph-9(0)*: 609.1±186.7 n=217; *kap-1(0)*: 1128.4±216.5 n=765; *kap-1(0); maph-9(0)*: 1128.4±280.4 n=673]; XBX-1::RFP [Control: 850.8±255.8 n=177; *maph-9(0)*: 973.4±282.0 n=203]. (E) Quantification of number of motors/second. KAP-1::GFP [Control: 0.107±0.022 n=14; *maph-9(0)*: 0.132±0.018 n=16]; OSM-3::GFP [Control: 0.102±0.25 n=16; *maph-9(0)*: 0.0649±.021 n=19; *kap-1(0):* 0.0773±.022 n=23; *kap-1(0); maph-9(0):* 0.0521±.016 n=20]; XBX-1::RFP [Control: 0.153±0.013 n=10; *maph-9(0)*: 0.150±0.023 n=11]. (F) Schematic summarizing the change in velocity and frequency of the indicated motors in *maph-9(0)* worms compared to control. (G) Localization of endogenously tagged MAPH-9 (green) and SPD-5, TBB-4 (magenta) in an adult male tail. Insets (right) display localization of MAPH-9 in cilia of an individual ray. Scale bar, 5 μm. (H) Mating efficiency: Quantification of percent of progeny from mating (cross-progeny) with control or *maph-9(0)* mutant hermaphrodites and males, as indicated. Values presented are mean±SD. p values are calculated by Welch’s t-test. Graphs present individual data points with horizontal bar representing the median.

We next studied the role of MAPH-9 on motor activity by directly measuring motor activity in *maph-9(0)* mutants. We measured the velocity of motor particles by generating kymographs of OSM-3 and KAP-1 movement in control and *maph-9(0)* mutant axonemes (Figure 3C). The anterograde movement of both KAP-1 and OSM-3 was significantly slower in *maph-9(0)* mutants compared to controls (Figure 3D), indicating that MAPH-9 promotes anterograde motor movement. We isolated OSM-3 motor movement by examining OSM-3 velocity in *kap-1(0)* mutants. Although, in this *kap-1(0)* mutant background, OSM-3 moved at the same speed in axonemes with and without MAPH-9 (Figure 3D), we found a lower frequency of OSM-3 anterograde tracks in *maph-9(0)* mutants compared to control (Figure 3E). In contrast, we found a higher frequency of KAP-1 tracks in the absence of MAPH-9 (Figure 3E). The change in frequency observed in *maph-9(0)* mutants to favor the slower kinesin-II and disfavor faster OSM-3 could explain the overall decreased velocity of the motors when they walk connected in IFT trains. In contrast to the anterograde motors, dynein component XBX-1 velocity was increased in *maph-9(0)* mutants (Figure 3D) and had no change in the frequency of retrograde tracks (Figure 3E). This ability of XBX-1 to walk faster and with the same processivity suggests that the deleterious effects we see on the velocity and processivity of anterograde motors is independent of the ultrastructural defects that arise in *maph-9(0)* mutants and instead likely reveals a direct role for MAPH-9 in modulating anterograde motor activity (Figure 3F).

Cilia in *C. elegans* sensory neurons mediate diverse interactions with the extracellular environment and a wide array of behaviors, for example the ability of male worms to locate and copulate with hermaphrodites (Lipton et al., 2004; Liu & Sternberg, 1995). To understand if MAPH-9 is required for cilia function, we assessed the mating ability of *maph-9(0)* mutants.

MAPH-9 localizes to all cilia of the male tail (Figure 3G), but, as in other sensory neurons, MAPH-9 was not required for ciliogenesis. Despite the presence of cilia, mating efficiency was significantly reduced in both male and hermaphrodite *maph-9(0)* mutants with the greatest observable reduction in mating efficiency when *maph-9(0)* mutant hermaphrodite and males were mated (Figure 3H). These data indicate that although MAPH-9 is not strictly required for ciliogenesis, MAPH-9 in the axoneme mediates cilia function.

### MAPH-9 localization is conserved in mammalian cells

Given the clear role of MAPH-9 in *C. elegans* cilia, we wanted to explore whether this function is conserved. MAP9 homologs are conserved in a large subset of metazoans, including in mice and humans. Although the MAP9 amino acid conservation is poor among metazoans, AlphaFold predictions of MAP9 (Jumper et al., 2021; Varadi et al., 2022) across species all contain a single ∼202-angstrom long alpha helix (Figure 4A-4B; Figure S3A-S3B; Table S1), which corresponds to the human MAP9 microtubule binding domain (Venoux et al., 2008). Initial characterization of MAP9/ASAP in U-2 OS cells found it to be localized to the mitotic spindle in dividing cells (Saffin et al., 2005), but did not explore MAP9 localization in interphase ciliated cells. We therefore validated a MAP9 antibody (Figure S4A-4E) to explore the localization of MAP9 in mammalian cells and tissues. Consistent with previous reports in U-2 OS cells (Saffin et al., 2005; Venoux et al., 2008), MAP9 localized to the mitotic spindle in RPE-1 cells (Figure S4F). In addition, MAP9 clearly localized to cilia in interphase RPE-1 cells (Figure 4C).

**Figure 4.**
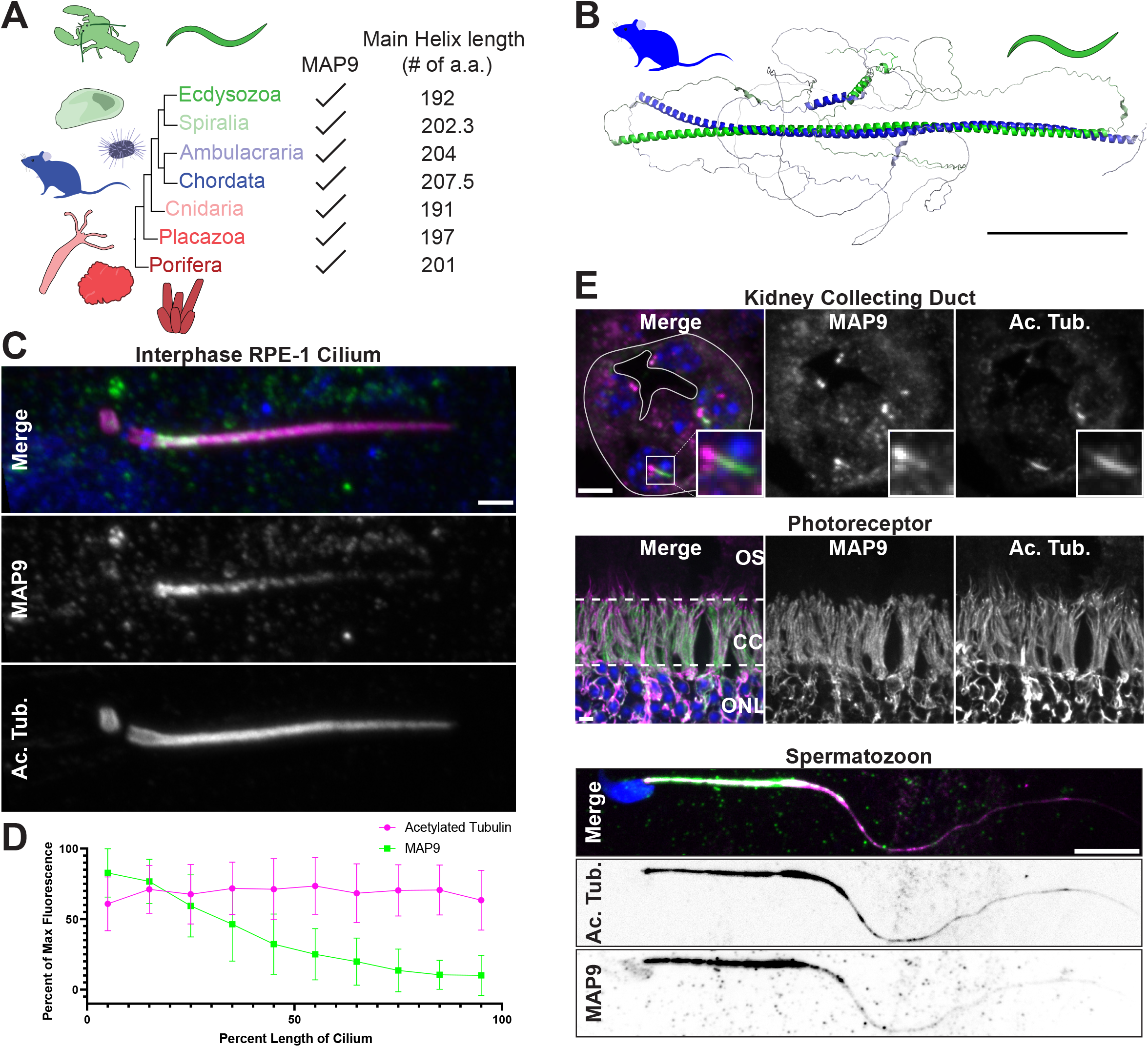
Localization of the MAPH-9 homolog MAP9 is conserved in mammalian cells. (A) MAP9 homologs with a consistent main alpha-helix length present throughout Metazoa. (B) Aligned AlphaFold structural predictions for mouse MAP9 (blue) and *C. elegans* MAPH-9 (green) Scale bar,100Å. (C) Localization of MAP9 (green), acetylated tubulin (Ac. Tub., magenta) and DNA (blue) in an isotropically expanded RPE-1 cell. Scale bar, 1 μm. (D) Percent of max fluorescence of background subtracted MAP9 and Ac. Tub. versus percent position down the length of the cilium. (E) Localization of MAP9 (green), Ac. Tub. (magenta), and DAPI (blue) in mouse tissues. Kidney collecting duct: Scale bar, 5 μm. Eye photoreceptor: outer segment (OS) connecting cilium (CC) outer nuclear layer (ONL). Scale bar, 5 μm. Spermatozoon: Scale bar, 10 μm.

Intriguingly, MAP9 localization in cilia was enriched in the proximal region adjacent to the centrosome and diminished toward the distal end of the axoneme (Figure 4C-4D). Although ultrastructural analysis of axonemes in RPE-1 cells is lacking, this localization pattern is consistent with the shorter microtubule doublet region found in mammalian MDCK-II cells (Kiesel et al., 2020). The localization of MAP9 to where doublets are typically found in mouse cilia (Deane et al., 2013; Gilliam et al., 2012; Konno et al., 2016) was also observed: MAP9 localized to the primary cilia of the collecting duct cells in the kidney; the connecting cilium in the photoreceptors of the eye; and to flagella in developing mouse spermatids and mature mouse spermatozoa (Figure 4E and Figure S4G-S4H). Together, these results indicate a conserved role for MAP9 in mammalian cilia, potentially in the doublet region of the axoneme.

## Discussion

Through endogenous labeling of MAPH-9, we found a remarkable preference of MAPH-9 for MTDs. This preference seems to in part be regulated by polyglutamylation, a post-translational modification that occurs on the C-terminal tails of α- and β-tubulin (Edde et al., 1990). Given the importance of steric suppression of these C-terminal tails to MTDs formation (Schmidt-Cernohorska et al., 2019), we speculate that MAPH-9 may contribute to B-tubule formation by helping to reduce the steric hinderance of tubulin C-terminal tails. This role is certainly in combination with other factors as: 1) MAPH-9 does not bind singlet microtubules *in vivo* and instead can discriminate MTDs as they are built; and 2) MAPH-9 depletion resulted in a loss of MTDs in only a subset of neurons. This regulation could be through specific association with tubulin isotypes, other tubulin posttranslational modifications, or association and/or competition with other MAPs. Further insight into the regulation of MAP9 and its physical interaction with MTDs would help our understanding of how MTDs are formed and stabilized.

MAPs have been proposed in many contexts to influence motor walking on cytoplasmic microtubules (Dixit et al., 2008; Ecklund et al., 2017; Lipka et al., 2016; Monroy et al., 2018; Monroy et al., 2020). The most prevalent phenotype we observed in *maph-9(0)* mutants was the deformation of the MTD structure. Given the damage molecular motors can impose on microtubules (Triclin et al., 2021), MAPH-9 might help to stabilize MTDs in the face of motor walking. Additionally, we found a role for MAPH-9 in anterograde motor movement in the axoneme, with a specific effect on the processivity of OSM-3. While two axonemal motors are not strictly required to build a functional cilium, this dual motor system is conserved among many metazoans that also have MAP9 homologs (Table S1), and perhaps MAP9 modulates these motors to allow for fine tuning of IFT speed and processivity (Scholey, 2013).

Finally, MAP9 has been posited as a target for cancer treatment given its localization to the mitotic spindle (Liu et al., 2018; Wang et al., 2020; Zhang et al., 2022); however, we have shown that MAP9/MAPH-9 has an important role in cilia in supporting MTDs and modulating motors. Further work understanding MAP9/MAPH-9 will give insight into how MTDs are formed as well as be a protein of interest in understanding ciliopathies.

## Acknowledgements

We thank Nina Peel, Guangshou Ou, and Kassandra Ori-McKenney for providing strains, protein, antibody, and advice. We also thank members of the Feldman lab for helpful discussions about the manuscript. This works was supported by CMB training grant T32GM007276 awarded to M.V.T.; NIGMS K99GM135489 awarded to M.D.S.; NIGMS K99GM131024 awarded to J.T.W.; R01GM136902 and R01GM133950 awarded to J.L.F.; R01NS082208 awarded to K.S; AHA Postdoctoral Fellowship 903408 awarded to S.R.W.; AHA Postdoctoral Fellowship 20POST35210767 awarded to J.W.F.; NIGMS of the National Institutes of Health under award numbers R35 GM130286 and R01 NS082208 awarded to T.S.; L.E.C. is a Damon Runyon Fellow supported by the Damon Runyon Cancer Research Foundation (DRG-2428-21). K.S. is a Howard Hughes Medical Institute Investigator. C.J. is supported by the Institut Curie, the French National Research Agency (ANR) award ANR-20-CE13-0011, and the Fondation pour la Recherche Medicale (FRM) grant DEQ20170336756. M.M.M. is supported by the Fondation Vaincre Alzheimer FR-16055p and the France Alzheimer grant 2023. M.G. is supported by Institut Curie 3-i PhD Program (IC-3i) and the European Molecular Biology Organization short-term fellowship 8843.This work was supported by Czech Science Foundation grant 19-27477X to Z.L. and institutional support from the CAS (RVO: 86652036) The project was supported by National Center for Research Resources (Stanford Imaging Award Number 1S10OD01227601). Some of the nematode strains used in this work were provided by the Caenorhabditis Genetic Center, which is funded by the NIH Office of Research Infrastructure Programs (P40 OD010440).

## Author Contributions

Conceptualization: M.V.T; J.M.; J.L.F.

Investigation – Mouse: J.W.F.; S.R.W.; M.V.T.; J.M.

Investigation – EM: R.D.F.

Investigation – Cell Culture: M.V.T.; J.T.W.

Investigation – PTM Tubulin: M.G.; M.M.M; C.J.

Investigation – *in vitro* MT binding: D.K.; L.Z.

Investigation – Mating: M.D.S.

Investigation – Phylogenetics: L.E.C.

Investigation – Strains: M.V.T.; J.M.; S.E.

Formal Analysis: M.V.T; J.M.; L.E.C.

Resources: J.L.F.; T.S.; K.S.; C.J.; L.Z.;

Writing – Original Draft: M.V.T; J.L.F.

Review & Editing: M.V.T; J.L.F.; J.M.

Visualization: M.V.T; J.M.; J.W.F. L.E.C. J.L.F.

Supervision: J.L.F.; J.M.

## Supplemental Material

### Supplemental Figure Legends

**Figure S1. MAPH-9 localization and preference for polyglutamylated tubulin; *maph-9(0)* mutant phenotypes**

(A) Whole adult worm endogenously expressing GFP::MAPH-9 with *glo-1* depleted by RNAi to reduce gut granule fluorescence. Scale bar, 50 μm.

(B) Localization of endogenously tagged MAPH-9 (green) and SPD-5 (magenta) through development. Top: Staged embryos with amphid neuron outlined and centriole at ciliary base (orange arrow) or cell body (gray arrow) indicated. Scale bar, 5 μm. Bottom: Magnified insets of centriole at ciliary base (orange boxes) or cell body (gray boxes) at different embryonic stages. Scale bar, 1 μm.

(C) Quantification of fluorescence intensity (a.u.) of MAPH-9 at the ciliary base vs cell body. Prebean: ciliary base; 342.3±2.52 n=3; cell body: 314.6±51.05 n=3. Bean: ciliary base: 633.56±147.2 n=10; cell body: 371.1±68.8 n=7. Comma: ciliary base: 1020±170.1 n=5; cell body: 356.8±36.8 n=5. 1.5 fold: ciliary base: 1571±623.6 n=9; cell body: 402.9±48.8 n=9.

(D) Quantification of fluorescence length (μm) of MAPH-9 and TBB-4 measured from the ciliary base marked by SPD-5. Time is measured in hours after 1.5-fold embryo stage. +1.5 hrs (550 mpf): 0.52±0.098 μm n=14. +2.5 hrs (610 mpf): 1.48±0.39 μm n=18. +3.5 hrs (670 mpf): 2.13±0.28 μm n=12. +4.5 hrs (730 mpf): 3.11±0.64 μm n=12. +5.5 hrs (790 mpf): 3.20±0.82 μm n=10. L1: 3.29±0.76 μm n=12.

(E) Total Internal Reflection Fluorescence images: Left and Middle: microtubules assembled from tubulin from wildtype (WT) mice (white arrowheads) or *Ttll1*^−/−^, *Ttll7*^−/−^, *Ttll1*^−/−^ and *Ttll7*^−/−^, or *Atat1*^−/−^ mice (orange arrowheads); Right: MAPH-9 localization. Scale bar, 5 μm.

(F) Schematic of endogenous *maph-9* locus with sgRNA cut sites and BFP replacement indicated to make *maph-9(0)* strain.

(G) Localization of TBA-5 in phasmid cilia. Scale bar, 1 μm.

(H) Length (μm) of TBA-5 localization from ciliary base. Control: 6.58±0.75 n=24. *maph-(0)*: 6.53±1.4 n=74; p=0.83. Values presented are mean±SD. P-value calculated by Welch’s t-test. Graphs present individual data points with horizontal bar representing the median.

(I) Localization of endogenously tagged TBB-4::GFP and SPD-5::GFP in indicated mutant backgrounds. For *kap-1(0); maph-9(0)* mutants, a representative image of two different mutant phenotypes are shown: Top: TBB-4 and SPD-5 localization length greater than 5 μm; Bottom; TBB-4 and SPD-5 localization length less than 5 μm. Scale bar, 1 μm

**Figure S2. Ultrastructure of amphids and ASI neuron in *maph-9(0)* mutants**

(A) Representative EM images of amphids neuron axonemes in control and *maph-9(0*) mutant worms. Scale bar, 200 nm. Numbered yellow arrowheads correspond to insets on right. Scale bar, 100 nm.

(B) Representative EM images of amphid neuron ASI axoneme in control and *maph-9(0)* mutant worms. Scale bar, 100 nm.

(C) Quantification of doublet, incomplete doublets, and singlet microtubules in ADL axonemes. Measurement at base (left) and then in sections in the middle segment going toward the distal segment (right).

**Figure S3. MAP9 phylogeny and predicted structure across species**

(A) Maximum likelihood tree from an alignment of the main alpha helix in MAP9 homologs with nodes indicating bootstrap support out of 1000 repeats. Species names are colored by group (right). Scale bar indicates amino acid substitutions per position.

(B) AlphaFold structural predictions for select species depicting well conserved main alpha helix. Accession number available in Table S1.

**Figure S4. MAP9 antibody validation and mammalian localization**

(A) Ponceau stained nitrocellulose containing indicated purified protein cut out for antibody preabsorption assay.

(B) Average fluorescence (a.u.) of MAP9 antibody staining over cilia length in RPE-1 cells with antibody mixture preabsorbed (PA) with expressed and purified GFP or GFP:MAP9. GFP preabsorbed: 79937±31484 n=19; GFP:MAP9 preabsorbed: 19177±3452 n=18; p<0.0001.

(C) Immunostained cilia from RPE-1 cells with antibody preabsorbed with expressed and purified GFP or GFP:MAP9. Scale bar, 1 μm.

(D) Western Blot of RPE-1 cells transfected to express GFP, GFP:MAP9, GFP:MAPH-9, probed with α-GFP (left) or α-MAP9 (right).

(E) Cilia from RPE-1 cells immunostained with Ac. Tub. (magenta) and MAP9 Invitrogen antibody (Left, green) (Monroy et al., 2020) and MAP9 ProteinTech antibody (Right, green). Scale bar, 1 μm.

(F) Mitotic RPE-1 cells immmunostained with antibodies against MAP9 (green in merge), Ac. Tub. (magenta in merge), Centrin, and DAPI. Scale bar, 5 μm.

(G) Elongating mouse spermatid immunostained with antibodies against MAP9 (green in merge) and α-tubulin (magenta in merge). Additional MAP9 localization to the manchette (“M”) is indicated. An overexposure of α-tubulin localization is shown at bottom to highlight the axoneme. Scale bar, 5 μm

(H) Step 8 round mouse spermatids (final stage before spermatid elongation using α-tubulin manchette) immunostained with antibodies against MAP9 (green in merge at left), Centrin (magenta in merge), and DAPI (blue in merge). Scale bar, 5 μm.

## STAR Methods

### *C. elegans* strains and maintenance

*C. elegans* strains were cultured at 20°C on plates of Nematode Growth Medium (NGM) covered with a lawn of OP50 *E. coli* (Sulston & Brenner, 1974). Strains used in this study are listed in the Key Resources Table.

### CRISPR/Cas9 cloning and editing

Genes were endogenously edited to create mutants or endogenously tagged proteins using the CRISPR Self Excising Cassette (SEC) method (Dickinson et al., 2015). Cas9 and appropriate sgRNAs were delivered using a plasmid based on pDD162, with the sgRNA (Key Resources Table) incorporated using the Q5 Site directed mutagenesis kit (New England BioLabs). Repair templates for each edit were generated by adding two homology arms flanking the appropriate SEC. Homology arms were generated by amplifying the region close to the desired insertion site (Key Resources Table). The repair vector was digested and homology arms were assembled using a HiFi DNA assembly reaction (New England BioLabs) accordingly to the manufacturer’s recommendation. DNA mixtures (sgRNA and Cas9 containing plasmid and repair template) were injected into the germline of adults worms, and CRISPR edited worms were selected by treatment with hygromycin followed by screening for appropriate expression and localization.

CRISPR edited worm lines were then backcrossed two times and homozygosed.

### RNAi

RNAi treatment was performed by feeding L4 worms with HT115 bacteria transformed with *glo-1* (RNAi) plasmid Bacteria were spread onto NGM plates supplemented with 1mM IPTG and 50 μg/mL Ampicillin and grown for 48 hours at room temperature away from light. L4 stage worms were left on RNAi plates at 25°C for 48 to 72 hours and their progeny were imaged.

### Mating efficiency assay

Three L4 hermaphrodites and three L4 males were placed on a 35mm NGM plate spread with OP50 *E. coli* and allowed to mate for 24 hours at 20°C. The males and hermaphrodites were then transferred to a fresh plate, allowed to mate for an additional 48 hours, and removed. All cross- and self-progeny were counted on a fluorescent dissecting scope with nuclear intestinal GFP from the males marking cross-progeny and the absence of GFP marking self-progeny. Each cross was performed in triplicate for each hermaphrodite/male genotype pair.

### Cell culture

RPE-1 cells were cultured at 37 °C in 5% CO_2_ in DMEM F12 (1:1) (Gibco) supplemented with Cosmic Calf Serum (HyClone). Cells were passaged with Trypsin 0.25% (Corning).

### Transfection

Cells were transfected with lipofectamine 3000 reagents (Invitrogen) according to the manufacturer’s directions.

### Mouse Tissue Handling

Wild-type adult mouse kidneys and eyes were dissected and fixed for 2 hours in 4% PFA in PBS rotating at 4°C. Tissue was washed two times in PBS and equilibrated in a graded series of sucrose solutions (10% and 20%). Tissue was flash frozen in O.C.T compound (Fisher Scientific) in a dry ice/95% ethanol slurry, and kidneys (coronal) and adult mouse eyes (transverse) were cryosectioned at 20µm. Cauda epididymal sperm were collected after extrusion from the epididymis in PBS. Developing spermatids were assessed via tubule squash preparations from mouse testes.

### Immunohistochemistry

#### Kidneys and eyes

Frozen sections were dried at room temperature, washed 1x in PBS (5 min. at room temperature), 1x in PBT (1% Triton X-100 in PBS), blocked (10% goat serum + 1% BSA in PBT) for 1 hour at room temperature. Samples were incubated with primary antibodies overnight at 4°C, washed 2x in PBT, incubated in species-specific secondary antibodies at room temperature for 1 hour, and then incubated for 5 mins with DAPI (4’,6-diamidino-2-phenylindole, Thermo Fisher Scientific) (0.5µg/mL in PBS).

#### Sperm

Epididymal sperm were attached to microscope slides for 30 min and fixed with 4% PFA in PBS for 15 min. The slides were washed with PBS three times before permeabilizing with -20°C methanol for 2 minutes. Slides were washed with PBS three times and incubated overnight at 4°C in antibody dilution buffer (ADB; 3% bovine serum albumin (BSA), 10% horse serum, 0.05% Triton X-100) containing primary antibodies. Slides were washed three times in PBS before applying secondary antibodies diluted in ADB and incubated overnight at 4°C. Secondary antibodies conjugated to Alexa 488, 568, or 633 against rabbit and mouse IgG (Thermo Fisher Scientific) were used at 1:500 dilution. Epididymal sperm immunofluorescence slides were mounted in Mowiol^®^ 4-88 (EMD Millipore) mounting medium containing DAPI (Thermo Fisher Scientific).

### In vitro Microtubules

Mouse wild-type tubulin as well as mouse tubulin with post-translation modification (*Ttll1*^*-/-*^, *Ttll7*^*- /-*^, *Ttll1*^*-/-*^*Ttll7*^*-/-*^, *Atat1*^*-/-*^) was purified as described in (Genova et. al., 2023).

*GMPCPP-lattice microtubules* (*GMPCPP* polymerized) with post-translation modification were polymerized from 4 mg/ml mouse tubulin (wild type, *Ttll1*^*-/-*^, *Ttll7*^*-/-*^, *Ttll1*^*-/-*^*Ttll7*^*-/-*^, *Atat1*^*-/-*^) for 1 h at 37°C in BRB80 supplemented with 2.7 mM *GMPCPP* (Jena Bioscience). The polymerized microtubules were centrifuged for 30 min at 18000 x g in a Microfuge 18 Centrifuge (Beckman Coulter). After centrifugation the pellet was resuspended in BRB80.

### In vitro Binding to Microtubule Lattice With Different Post-Translation Modification

Mouse *GMPCPP*-wild-type microtubules were immobilized on anti-tubulin antibody treated channels (Siahaan et al., 2022). Then the channels were flushed with 40 μl of BRB80 to remove unbound microtubules and a snapshot was taken to mark the position of wild type microtubules. Subsequently *GMPCPP*-mouse microtubules with post-translation modification (*Ttll1*^*-/-*^,*Ttll7*^*-/-*^, *Ttll1*^*-/-*^*Ttll7*^*-/-*^, *Atat1*^*-/-*^) were immobilized and the channels were flushed with 40 μl of BRB80 to remove unbound microtubules and again a snapshot was taken to mark the position of microtubules with post-translation modification. Finally, the flow chamber was flushed with 10 μl of 20 nM GFP-MAPH9 diluted in the assay buffer (BRB80, 0.2% Tween20, 0.5 mg/ml Casein, 20 mM D-glucose, 0.22 mg/ml glucose oxidase and 20 mg/ml catalase). Time-lapse image sequences of MAPH9 binding to microtubules were recorded for 1 min at the rate of 1 frame per second with an exposure time of 100 ms. Data from three independent experiments was collected, each experiment was repeated at least on three days.

### Microscopy

#### C. elegans Imaging

Worms were mounted on an agarose pad (5% agarose dissolved in M9) immersed in 1mM levamisole (Sigma-Aldrich) sandwiched between a microscope slide and no. 1.5 coverslip. Embryos were mounted the same way without levamisole.

#### TIRF microscopy

Total internal reflection fluorescence (TIRF) microscopy experiments were performed on an inverted microscope (Nikon-Ti E, Nikon-Ti2 E) equipped with 60x or 100x NA 1.49 oil immersion objectives (Apo TIRF or SR Apo TIRF, respectively, Nikon Instruments) and either Orca Flash 4.0 sCMOS (Hamamatsu) or PRIME BSI (Teledyne Photometrics) cameras. An additional 1.5x magnifying tube lens was used. Microtubules were visualized by using an epifluorescence lamp and GFP-MAPH9 with HiLyte-647 tubulin (Siahaan et al., 2022)were visualized sequentially by switching between microscope filter cubes for Cy5 and GFP channels. The imaging setup was controlled by NIS Elements software (Nikon Instruments).

Flow chambers for TIRF imaging assays were prepared as described previously (Fink et al., 2009). Channels were treated with anti-biotin antibody solution (Sigma-Aldrich, 1 mg/ml in PBS) or alternatively with anti-tubulin antibody (Sigma-Aldrich, 1 mg/ml in PBS) solution after 5 minutes, followed by one-hour incubation with 1% Pluronic F127 (Sigma-Aldrich).

#### Dye Filling Assay

Young adult worms were incubated in 5 μg/ml DiI (1,1′-dioctadecyl-3,3,3′, 3′- tetramethylindocarbocyanine perchlorate, Medchemexpress) diluted in M9 for 1hr. Worms were then washed 3 times in M9 and placed onto NGM plates for 2 hrs for destaining. Dye filled worms were mounted and imaged as above.

#### Late-Stage Embryo Imaging

Embryos at the 1.5 fold stage were selected by manual picking and then excised from the NGM plate to a humidified chamber to be aged. At the designated time after the 1.5 fold stage, embryos were placed in an airtight chamber with an intake valve for CO_2_ delivery from dry ice and an exhaust valve. After incubation with CO_2_ for 15 minutes, the chamber was sealed and imaged as described below.

#### Spinning-disk confocal microscopy

Unless indicated, all images were acquired using a Nikon Ti-E inverted microscope (Nikon Instruments), a Yokogawa CSU-X1 confocal spinning disk head, and an Andor Ixon Ultra back thinned EM-CCD camera (Oxford Instruments - Andor), controlled by NIS Elements (Nikon Instruments). Images were obtained using a 60x (NA= 1.4) or 100x Oil Plan Apochromat objective (NA= 1.45). Z stacks were acquired using a 0.2 μm step. For timelapse, images were acquired at 150 ms exposure with no delay for 1 minute.

#### Structured illumination microscopy (SIM)

Structured illumination microscopy images were acquired on an OMX BLAZE V4 microscope (GE healthcare) in the Stanford Cell Science Imaging Facility equipped with a U-PLANAPO 100x SIM (NA = 1.4) objective (Nikon Instruments) and Evolve 512 emCCD cameras (Teledyne Photometrics) controlled by DeltaVision software. Structured Illumination images were generated using SoftWoRx (GE healthcare). Images were adjusted for brightness and contrast using ImageJ.

#### Ultrastructure expansion microscopy (U-ExM)

RPE-1 cells were grown on 12 mm, #1.5 glass coverslips. Coverslips were fixed in methanol at 20°C for 10 minutes and washed with 1X PBS. Coverslips were incubated overnight at 37°C in an acrylamide/formaldehyde solution (AA/FA, 0.7% formaldehyde, 1% acrylamide in PBS).

Gelation was allowed to proceed in monomer solution (19% sodium acrylate, 10% acrylamide, 0.1% bis-acrylamide, 0.5% ammonium persulfate-APS, 0.5% TEMED) and the coverslips were discarded. Gels were boiled at 95°C in denaturation buffer (200 mM SDS, 200 mM NaCl, 50 mM Tris pH 9) for 1 hour. Denaturation buffer was removed, gels were washed with multiple water rinses and allowed to expand in water at room temperature overnight. Small circles (approximately 5 mm in diameter of each expanded gel) were excised and incubated with primary antibodies diluted in PBSBT buffer (3% BSA, 0.1%Triton X-100 in PBS) on a nutator at 4°C overnight. The next day, gels were washed three times with PBSBT buffer and incubated with secondary antibodies and 5 μg/mL DAPI diluted in PBSBT, on a nutator at 4°C overnight.

Gels were washed once with 1X PBS and three times with water, and placed in a glass-bottom, poly-L-lysine treated 35mm plate to image.

#### Electron microscopy

Age-matched *maph-9(0)* mutants and wild-type worms were prepared for conventional electron microscopy by high-pressure freezing and freeze-substitution. Worms in M9 media supplemented with 20% BSA and OP50 *E. coli* were put in specimen carriers with a 50μm deep well (Technotrade International), covered with the flat side of a type B specimen carrier (Technotrade International), and frozen using a Leica EM ICE high pressure freezer. Freeze-substitution was done with a Leica AFS2 unit in acetone with 1% OsO_4_, 0.1% uranyl acetate, 1% methanol and 3% water (Buser & Walther, 2008; Walther & Ziegler, 2002). After substitution, samples were rinsed in acetone, infiltrated, and then polymerized in Eponate 12 resin (Ted Pella). Serial 50-nm cross-sections through the amphid and phasmid cilia of worms were cut with a Leica UC7 ultramicrotome using a Diatome diamond knife. Sections were picked up on Pioloform-coated slot grids and stained with uranyl acetate and Sato’s lead (Sato, 1968). Sections were imaged with an FEI Tecnai T12 transmission electron microscope at 120 kV using a Gatan Rio 4k × 4k camera and DigitalMicrograph.

### Image Quantification

#### B-tubule circularity measurement

Measurements were performed using the Fiji package of the ImageJ2 software (2.9.0) on electron microscopy images of both left and right phasmid pairs at different position along the length of the middle segment. B-tubules were manually traced with the freehand tool and the roundness value was determined by analyzing the trace for circularity (4π(area/perimeter^2^).

#### Motor Tracking

Time lapse imaging was imported into image J using the KymographClear2.0 plugin and kymographs were generated. To calculate speeds, tracks were drawn manually and imported into KymographDirect software.

#### In vitro MAPH-9 density estimation

MAPH9 density on the microtubules with different post-translation modification was measured In FIJI by drawing a rectangle around the microtubule and measuring the average intensity per pixel. The rectangle was then moved to an area directly adjacent to the microtubule where no microtubule is present and the average intensity per pixel was measured again and subtracted from of the average intensity per pixel the microtubule. The final density of MAPH-9 was normalized to the average of density MAPH-9 on mouse wild type microtubules.

### Protein Purification

Protein expression plasmids were transformed into BL21 DE3 cells (Sigma-Aldrich). Starter cultures were innoculated with one colony and grown overnight in 5mL of LB at 37°C supplemented with kanamycin. The starter culture was added to 500 mL of LB with kanamycin, grown at 37°C until OD600 was 0.6, then grown at 18°C for 30 minutes and induced with a final concentration of 100 µM IPTG. Bacteria were pelleted and resuspended in lysis buffer (250 mM NaH_2_PO_4_, pH 8.0, 2.5 M NaCl, Halt Protease Inhibitor Cocktail (Thermo Fisher Scientific)), lysed using Cell Disruptor MC (Constant System), and centrifuged at 3000 x g for 10 minutes. The supernatant was added to nickel resin Ni-NTA Agarose (Invitrogen) equilibrated in lysis buffer and incubated on a nutator for 1 hour at 4°C. The agarose was centrifuged at 800 x g for one minute and washed with wash buffer (250 mM NaH_2_PO_4_, pH 8.0, 2.5 M NaCl, 20 mM imidazole, pH 6.0) 3 times. Protein was eluted with elution buffer (250 mM NaH_2_PO_4_, pH 8.0, 2.5 M NaCl, 250 mM imidazole, pH 6.0).

### Antibody Preabsorption

Proteins were resolved by SDS-PAGE and transferred to nitrocellulose membrane. The membrane was stained with Ponceau S solution (Sigma-Aldrich) for 15 minutes and rinsed twice with water. The membrane around the bands corresponding to GFP and GFP::MAP9 was excised. The membranes were blocked with 5% milk in TBST buffer (Tris-buffered saline with 0.1% Tween-20) for 1 hour at room temperature rinsed twice in TBST buffer, and twice in PBSBT buffer (3% BSA, 0.1% Triton X-100 in PBS), and then incubated with primary antibody mixture with MAP9, and acetylated tubulin antibodies on a nutator for 5 hours at room temperature. The membranes were then discarded and the antibody mixture was added onto coverslips to stain them for immunofluorescence as described above.

### Western Blotting

Samples were denatured with 6X sodium dodecyl sulfate (SDS) Laemmli buffer and boiled at 95°C for 10 minutes. Proteins were separated by SDS-PAGE and transferred to a nitrocellulose membrane (Bio-Rad). The membrane was blocked with nonfat milk at room temperature for 1 hour and then incubated with primary antibodies at 4°C overnight. The membrane was incubated with secondary antibodies at room temperature for 1 hour. All blots were imaged with an Azure 600 (Azure Biosystems) and analyzed using image J.

### Phylogenetics and AlphaFold structural predictions

Homologs to human MAP9 (see Table S1) were identified through: 1) BLAST or PSI-BLAST to NCBI databases using default values 2) proteins containing MAP9 domain IPR026106 that also contained AlphaFold v2 structural predictions (Jumper et al., 2021; Varadi et al., 2022). Protein sequences of homologs were aligned using Muscle (Edgar, 2004) and trimmed to conserved parts (total of 92 positions) within the main alpha helix in JalView (Waterhouse et al., 2009). A maximum likelihood tree was constructed using PhyML 3.0 (Guindon et al., 2010) with bootstrap support values from 1000 repeats and substitution model chosen by the Akaike Information Criterion (Lefort et al., 2017). Nodes with support less than 250/1000 are not shown on tree after visualization in FigTree v1.4.4. AlphaFold v2 structural prediction images and the identity of the longest right-handed alpha helix were obtained from the AlphaFold Protein Structure database and associated mmCIF files. Mouse and worm MAP9 homologs were aligned via pymol super and colored from grey to green/blue by pLDDT (spectrum b, grey lightblue blue, AF-Q3TRR0-F1-mod; spectrum b, grey palegreen green, AF-Q18452-F1-mod).

### Ethics statement

All mice were bred at Stanford University (Stanford, CA) or purchased from Jackson Laboratory (JAX) in accordance with the National Institutes of Health and U. S. Department of Agriculture criteria, and protocols for their care and use were approved by the Institutional Animal Care and Use Committees (IACUC) of Stanford and JAX.

