## Supplemental Figures for "MAP9/MAPH-9 supports axonemal microtubule doublets and modulates motor movement"

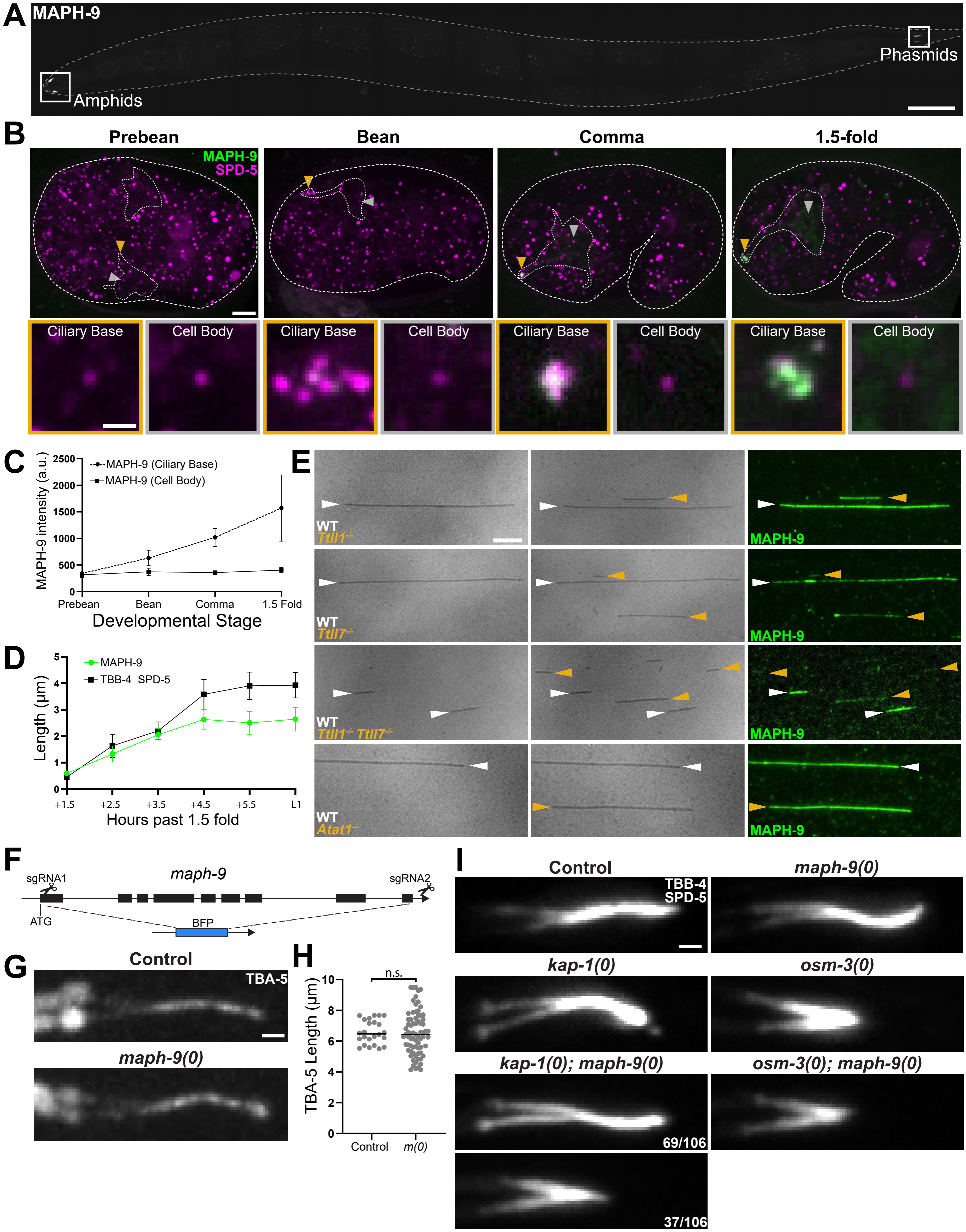


**Figure S1. MAPH-9 localization and preference for polyglutamylated tubulin; *maph-9(0)* mutant phenotypes**

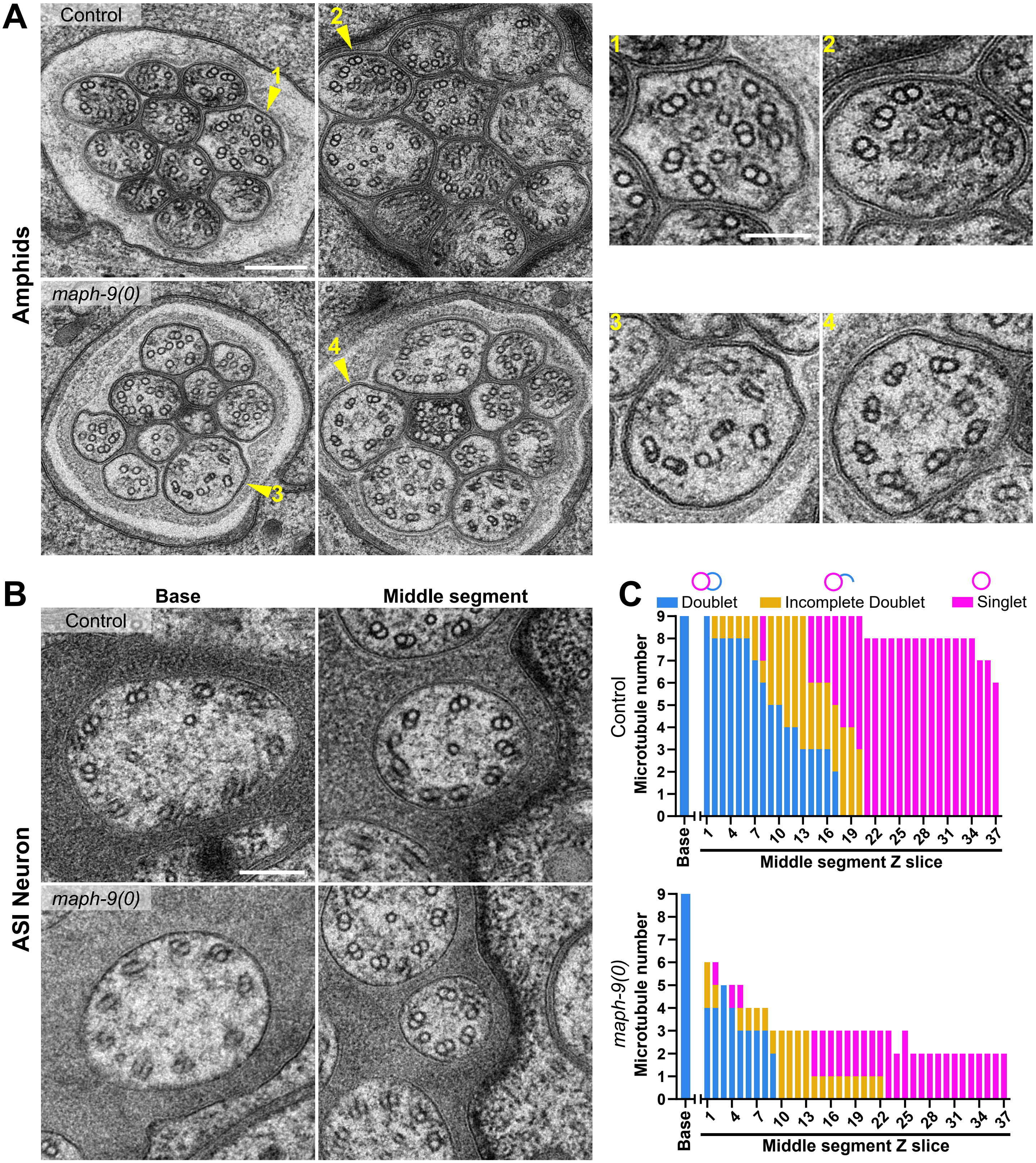


**Figure S2. Ultrastructure of amphids and ASI neuron in *maph-9(0) mutants***

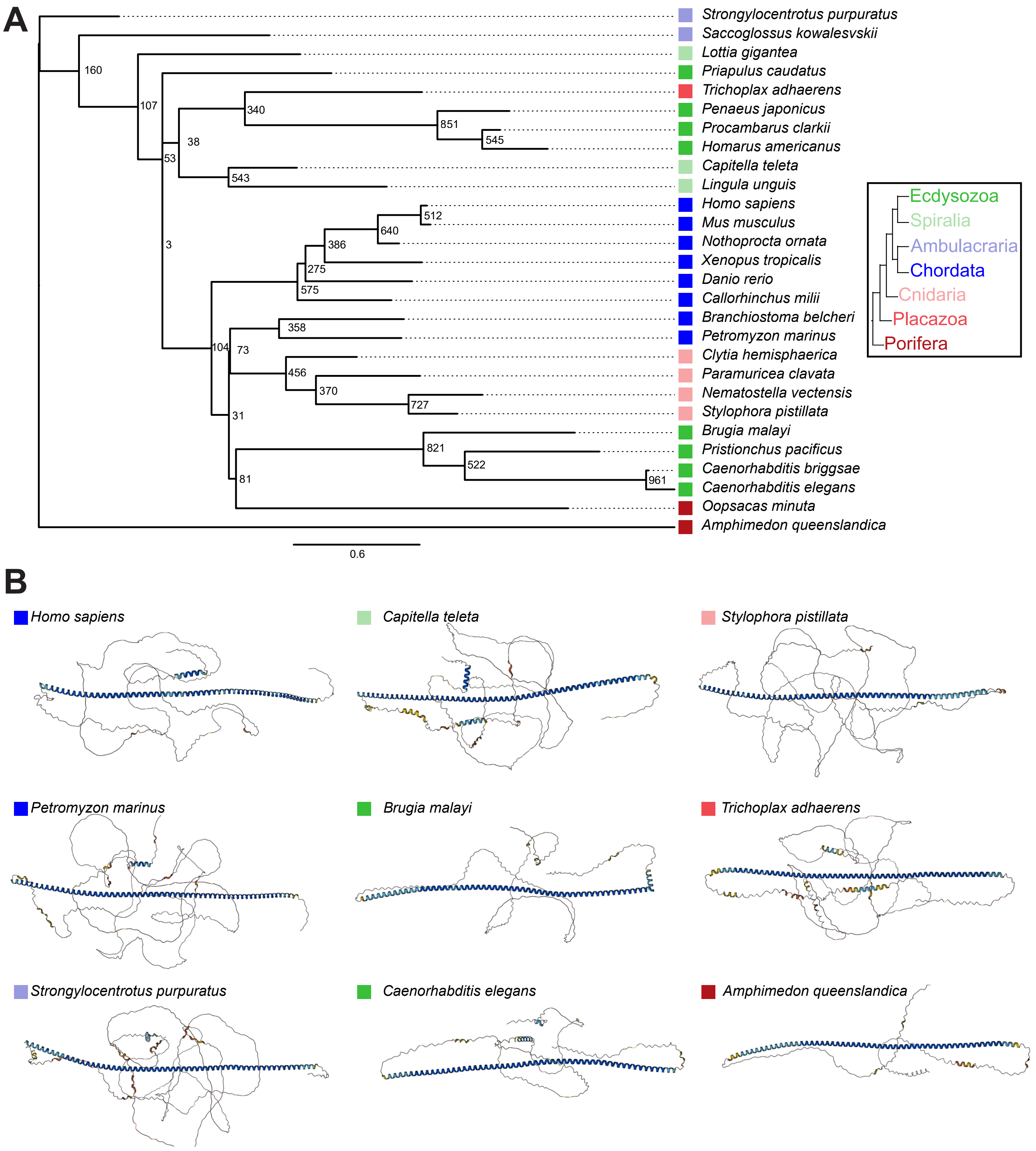


**Figure S3. MAP9 phylogeny and predicted structure across species**

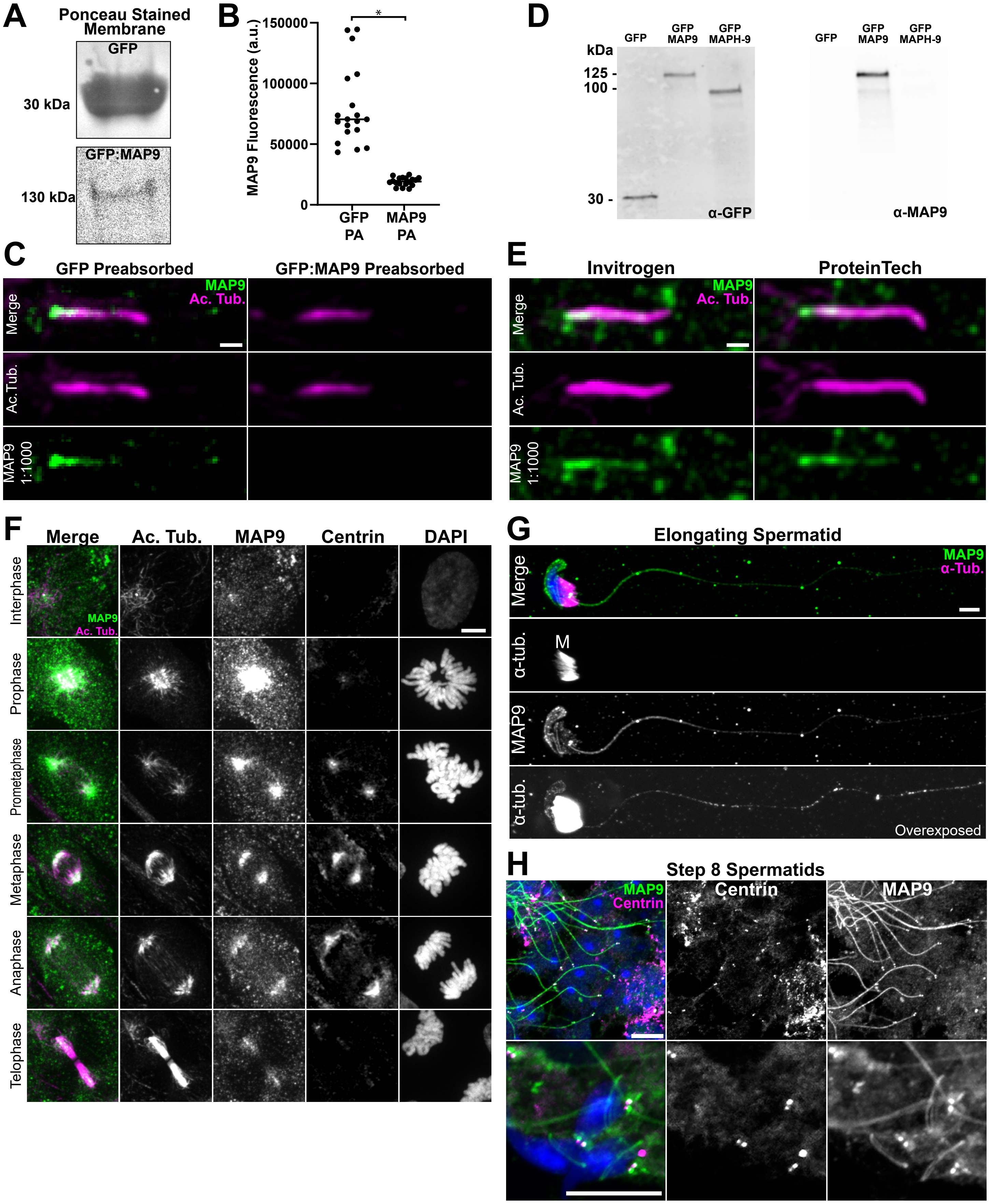


**Figure S4. MAP9 antibody validation and mammalian localization**
